# Alcohol Use Disorder and Smoking-Associated Molecular Alterations in Human Prefrontal Cortex in Single Nucleus (sn) RNA-Seq

**DOI:** 10.64898/2026.09.24.753990

**Authors:** Arpita Joshi, Pietro Paolo Sanna

## Abstract

Chronic smoking worsens alcohol-related brain injury and impairs neurocognitive recovery, yet the cell-type-specific molecular interactions between alcohol use disorder (AUD) and smoking in the human brain remain largely unexplored. We analyzed single-nucleus RNA-seq from the prefrontal cortex of 73 individuals (614,932 nuclei, 24 cell types), comparing AUD versus controls, smoking versus controls, and combined AUD+smoking versus controls. In excitatory neurons, neuroinflammatory, complement, GPCR, kinase, proteostatic, and extracellular-matrix pathways confined to one or two subtypes in either condition alone expanded across nearly all subtypes when AUD and smoking co-occurred. Inhibitory neurons showed a parallel but distinct expansion, additionally recruiting translational regulation, RNA splicing, and deubiquitination pathways. Smoking alone produced widespread repression of bioenergetic, proteostatic, and synaptic pathways in excitatory neurons, whereas AUD produced mixed pathway activation and repression in excitatory neurons and broad inflammatory activation in inhibitory neurons. In the combined condition, microglia exhibited paradoxical repression of phagocytosis, TYROBP signaling, and translational machinery, consistent with an exhaustion-like state, accompanied by an apparently compensatory shift in inflammatory signaling toward astrocytes, oligodendrocyte precursor cells, and oligodendrocytes, which showed coordinated inflammatory activation. Vascular cells showed an endothelial–VLMC dichotomy, with endothelial repression of RNA processing and proteostasis contrasting with perivascular activation. Our analysis identifies bioenergetic and proteostatic stress as the predominant signature of smoking, whereas immune and metabolic activation characterize AUD. In AUD+Smoking, these alterations extend across more neuronal subtypes and include neuroinflammation, complement activation, altered signaling, proteostatic stress, and post-transcriptional remodeling associated with greater cortical dysfunction in co-occurring AUD and smoking.

## INTRODUCTION

Alcohol Use Disorder (AUD) is a chronic, relapsing brain disorder characterized by an impaired ability to stop or control alcohol use despite adverse social, health, or occupational consequences [1], affecting approximately 28 million adults in the United States alone [2]. Daily tobacco use is a major public health concern linked to numerous diseases, with global prevalence estimates indicating over 1 billion smokers worldwide [3]. Notably, there is a high comorbidity between AUD and smoking, with studies showing that up to 80% of individuals with AUD also smoke regularly, and conversely, about 30% of smokers meet criteria for AUD, far exceeding rates in the general population [4–6]. This overlap is not merely coincidental; genetic, environmental, and neurobiological factors contribute to the co-occurrence, potentially exacerbating health risks through molecular mechanisms [7–9].

At the cellular level, chronic alcohol exposure in AUD induces widespread dysregulation, including mitochondrial dysfunction, impaired protein and lipid homeostasis, oxidative stress, and neuroinflammation, particularly in brain regions like the PFC and amygdala [10–13]. These changes can lead to altered neuronal plasticity, glial activation, and immune cell responses, with microglia playing a key role in driving AUD pathology through heightened inflammatory signaling [14, 15]. Similarly, chronic daily smoking exerts profound cellular effects, such as chronic inflammation, and DNA damage, [16–19]. Co-occurrence of smoking is associated with additive effects on health outcomes in AUD, including adversely affecting cognition [20–22], increased cancer incidence [23, 24], and increased mortality risk [25–28]. Comorbid chronic cigarette smoking also impairs neurobiological and cognitive recovery in abstinent alcoholics [29]. Neuroimaging studies further associate concurrent smoking with greater regional cortical metabolic abnormalities and gray-matter loss in individuals with AUD and impaired recovery of cortical perfusion during abstinence [30–32].

Traditional bulk tissue analyses have provided valuable insights into these conditions but fail to reveal cell-type-specific heterogeneity. Single-cell and single-nucleus RNA sequencing (scRNA-seq/snRNA-seq) offer a powerful approach to dissect these effects at cellular resolution, delineating regulatory perturbations in AUD involving nucleus accumbens, medium spiny neurons, cortical excitatory and inhibitory neuronal populations, oligodendrocytes, microglia, and other glial cell types.” [33, 34]. scRNA-seq transcriptomic studies of tobacco exposure identified immune, inflammatory, vascular, and epithelial regulatory perturbations involving neutrophils, cytotoxic T cells, vascular lesion–associated inflammatory programs, and bronchial epithelial cell states [35–37]. Single-nucleus studies in rodents identified nicotine/tobacco-associated regulatory remodeling in ventral tegmental dopaminergic and non-dopaminergic neurons, astrocytes, oligodendrocytes, and microglia [38]; as well as developmental perturbations in excitatory and inhibitory neuronal populations following prenatal e-cigarette exposure [39]. Bulk transcriptomic studies of human postmortem brain tissue identified convergent alcohol- and smoking-associated regulatory perturbations in the prefrontal cortex (PFC) and ventral tegmental area, including alterations in glutamatergic signaling, neuroplasticity, myelination-associated pathways, and stress-related molecular programs [40, 41]. Despite these advances, the effects of combined AUD and smoking on the central nervous system (CNS) at the single-cell level which could uncover interacting pathways in comorbid individuals, remain largely unexplored.

Here, we examined cell-type-specific transcriptional signatures in the PFC associated with AUD, daily smoking, and their co-occurrence to identify potential additive interactions relevant to the greater cortical dysfunction observed in comorbid individuals, using data published by Warden et. al [34]. The PFC is involved in decision-making and loss of control over drug intake and individual differences in prefrontal functioning have been implicated in vulnerability to substance abuse [42]. The present analyses show that smoking and AUD converge to produce a broader cortical pathology than either condition alone. Our analysis identifies bioenergetic and proteostatic stress as the predominant signature of smoking, whereas immune and metabolic activation characterize AUD. In the combined condition, these alterations extend across more neuronal subtypes and include neuroinflammation, complement activation, altered signaling, proteostatic stress, and post-transcriptional remodeling associated with greater cortical dysfunction in co-occurring AUD and smoking.

## MATERIALS AND METHODS

### Cell-type Annotations and Quality Control

The raw feature-barcode-matrix files for each sample from GEO (gene expression omnibus) were converted into a universal data object combining the separate metadata for each sample in the form of a compressed h5ad [43] file. We created new cell-type annotations by using the human neo-cortex cell-type labels as a reference using Allen Institute’s brain cell atlas, MapMyCells [44] web-tool. Quality control (QC) was performed to eliminate noisy cells by setting a minimum threshold of non-zero counts in at least 100 genes, thereafter, only protein-coding genes were retained, filtered using GENCODE version 44 annotations [45], the UMAP embedding in Figure-1 (a) shows the segregation of cell-type clusters at this QC stage. This was followed by correcting for ambient RNA by modelling it using scAR (single cell ambient remover) [46], which uses probabilistic deep learning with the underlying assumption that ambient signals introduced during sample preparation are drawn from a probability distribution with fixed probabilities for each feature, Figure-1 (b) shows the UMAP embedding after ambient RNA removal which does lead to relatively sharper projections of the cell-type clusters. Finally, the doublets were removed from the dataset using the SOLO model [47] which combines a variational autoencoder with an additional neural network layer to classify doublets based on learned embeddings, Figure-1 (c) shows the final UMAP embedding after all the QC steps.

**Figure-1:**
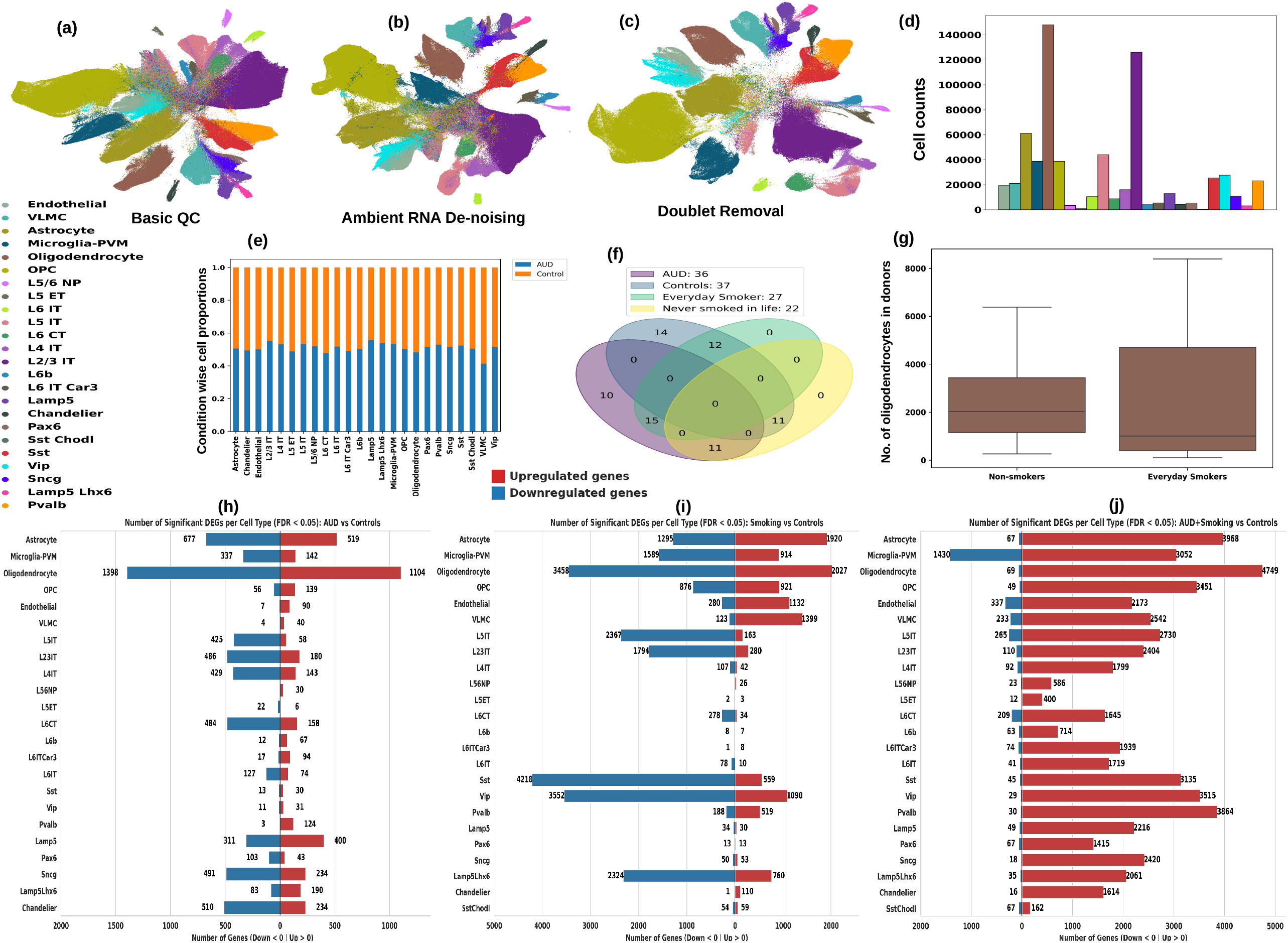
Summary of single-cell RNA-seq data with 73 AUD and Control samples from [34]. (a) UMAP embedding showing cell-type clusters past basic quality control (QC) of filtering cells and genes based on thresholding of gene and cell counts; (b) UMAP embedding showing the cell-type clusters after ambient RNA removal; (c) UMAP embedding showing cell-type clusters after doublet removal. Each QC stage increases the cluster separation especially among similar cell-types. (d) Number of cells in each cell-type establishing the prevalence of oligodendrocytes and L2/3 IT excitatory neurons. (e) Proportions of each cell-type in the AUD and Control categories. (f) Overlap of samples in the four categories analyzed in this work, namely, AUD, Control, Everyday Smokers and Non-Smokers. (g) Box-plot depicting the proportion of oligodendrocytes in everyday smokers and non-smokes (without AUD in each instance); oligodendrocytes in non AUD smokers versus non-smokers were the only cell-type category significantly depleted in the smoking population, no other cell-type population was depleted in any of the comparisons. (h, i, j) Number of most significant Differentially Expressed Genes (DEGs) per cell-type in each of the three comparisons. (h) AUD vs Controls; (i) Smoking vsControls; (j) AUD+Smoking vs Controls.

### scVI and Extensions for Compositional and Differential Analysis

snRNA-seq data were analyzed using scVI (single cell variational inference) [48, 49] version 1.0.4, a deep-learning based tool for single data analysis. Leveraging PyTorch [50] and AnnData [51] frameworks, scVI uses hierarchical Bayesian modeling [52] to map cell-by-gene expression data into a latent space, supporting downstream tasks such as clustering, visualization, and differential expression analysis. The model operates as a variational autoencoder [53] capturing gene expression patterns across cells with a zero-inflated negative binomial (ZINB) distribution. The data was corrected for age, sex and sequencing site of individual samples. The scVI model was trained with default hyperparameters: with at least 1000 epochs with an early stopping patience of 500, for training and validation error convergence.

### Other Bioinformatics Tools and Methods

We used Python version 3.11.4 and R version 4.3.0. For pathway analysis we used Python’s implementation of Gene Set Enrichment Analysis (GSEA) [54], gseapy, version 1.0.6, with Gene Ontology (Biological Process 2023 (GO: BP), Cellular Component 2023 (GO:CC), Molecular Function 2023 (GO: MF), ‘WikiPathway_2023_Human’ ‘MSigDB_Hallmark_2020’, ‘KEGG_2021_Human’ collections; pathways with FDR q-val <= 0.05 were considered significant.

## RESULTS

### Transcriptional Atlas of snRNA-seq Data

We used snRNA-seq data from the PFC of 73 individuals, 36 classified as AUD and 37 classified as control samples from the pre-frontal cortex. Amongst these samples, to identify smokers and controls we respectively segregated individuals who were reported to be smoking everyday/7 days a week and the ones that reported never having smoked in life, the overlap of the various categories is delineated in Figure-1 (f). After filtering out low quality cells (in terms of a threshold of gene counts), removing doublets using SOLO [47], and ambient RNA removal using scAR [55], a total of 614,932 nuclei remained, Figure-1 (a) shows the UMAP embedding of the entire dataset and Figure-1 (d) shows their cell-type composition and Figure-1 (e) shows the distribution of nuclei in various cell-types in the AUD and Control categories. Based on marker gene analysis, Leiden clustering and previously published data [56, 57], 24 cell-types were annotated in 7 major groups. Excitatory neurons comprised 195,926 nuclei (31.9%) and included L2-3 IT, L4 IT, L5 IT, L5 ET, L5-6 NP, L6 CT, L6 IT, L6 IT Car3, and L6b populations. Inhibitory neurons comprised 105,678 nuclei (17.2%) and included Chandelier, Lamp5, Lamp5 Lhx6, Pax6, Pvalb, Sncg, Sst, Sst Chodl, and Vip subtypes. Non-neuronal populations included oligodendrocytes (142,857; 23.2%), astrocytes (59,217; 9.6%), microglia (38,588; 6.3%), OPCs (37,947; 6.2%), VLMCs (18,877; 3.1%), and endothelial cells (15,842; 2.6%). Summary of cell-type abundance and distribution across AUD and control samples can be found in Figure-1 (d) and (e).

### Compositional Analysis using scCODA

To test the compositional changes in various cell-types among the three comparisons, namely, (1) AUD versus Controls (both non-smoking), (2) Everyday Smokers versus Non-smokers (both without AUD), and (3) Everyday Smokers with AUD versus Non-smoking Controls, we used the single-cell compositional data analysis (scCODA) framework. scCODA is a Bayesian model for detecting differential cell-type abundances in scRNA-seq data, tackling compositionality biases and low replicates by jointly modeling proportions via a Dirichlet-Multinomial likelihood with a log-link function for covariates. It fixes one reference cell type for relative changes, uses a logit-normal spike-and-slab prior for sparse variable selection, and infers posteriors via Hamiltonian Monte Carlo sampling. We only found differential compositional changes significantly perturbed in non-AUD smokers vs non-smokers among oligodendrocytes, which were reduced in everyday smokers; Figure-1 (g) shows the boxplot depicting the distribution of cell counts in the two categories, the mean counts for everyday smokers is significantly lower.

### Differential Analysis of Gene Expression

We compared levels of gene expression in cells isolated from individuals with AUD, individuals that were Everyday Smokers, and individuals that were Everyday Smokers and also had AUD, versus the Control samples by cell-type and identified in all 10,005 unique differentially expressed genes (DEGs) (filtered only by FDR corrected p-value at a threshold of <0.05). With only FDR filtering, in Figure-1 (h), (i), and (j) respectively, we show the distribution of the number of DEGs by cell-type for the three comparisons: AUD vs Controls, Smoking vs Controls and AUD+Smoking vs Controls. In order to segregate the more potentially biologically meaningful DEGs, we performed a stringent filtering by removing DEGs that were expressed in less than 10% of the corresponding cell-type and additionally were expressed at a level greater than the mean gene expression level of the corresponding cell-type, yielding 139 DEGs in AUD vs Controls comparison, 302 DEGs in Smoking vs Controls comparison, and 448 DEGs in AUD+Smoking vs Controls comparison. Finally, we intersected these three DEG lists to show the cell-type specific perturbation of these common 104 DEGs in all the three comparisons in Supplementary Figure-1. In AUD vs Controls and Smoking vs Controls comparison, most genes showed a strong signature of repression, with 70.3% (of which Oligodendrocytes have 40.9%) in prior and 84.4% (of which Sst inhibitory neurons have 58.2%, followed by Vip interneurons at 49% and Oligodendrocytes at 47.7%) in the latter. On the other hand, in AUD+Smoking vs Controls comparison, only 26.2% DEGs were repressed and the repressed DEGs are mostly from microglial cells (which have 58.1% of the repressed DEGs in this comparison). Most DEGs in this comparison (including the ones downregulated in some cell-types) show strong upregulation (Figure-1 (j)).

### Excitatory Neurons’ Responses in AUD with Smoking

GSEA across excitatory neuron subtypes revealed marked cell-type expansion of pathway dysregulation when AUD and smoking co-occurred (Figure-2). Pathways enriched in only one or a few subtypes in AUD or Smoking alone became engaged across most or nearly all excitatory populations in AUD+Smoking. This expansion was especially prominent for neuroinflammatory and immune signaling. TNF-alpha signaling via NF-kB and the TYROBP causal network were limited mainly to L2-3 IT and selected deep-layer populations in the single conditions but showed positive NES across nearly all excitatory subtypes in the combined condition. Inflammatory response, interferon-gamma response, cytokine-cytokine receptor interaction, cytokine activity, cytokine-receptor activity, and defense-response pathways showed the same broader distribution.

**Figure-2:**
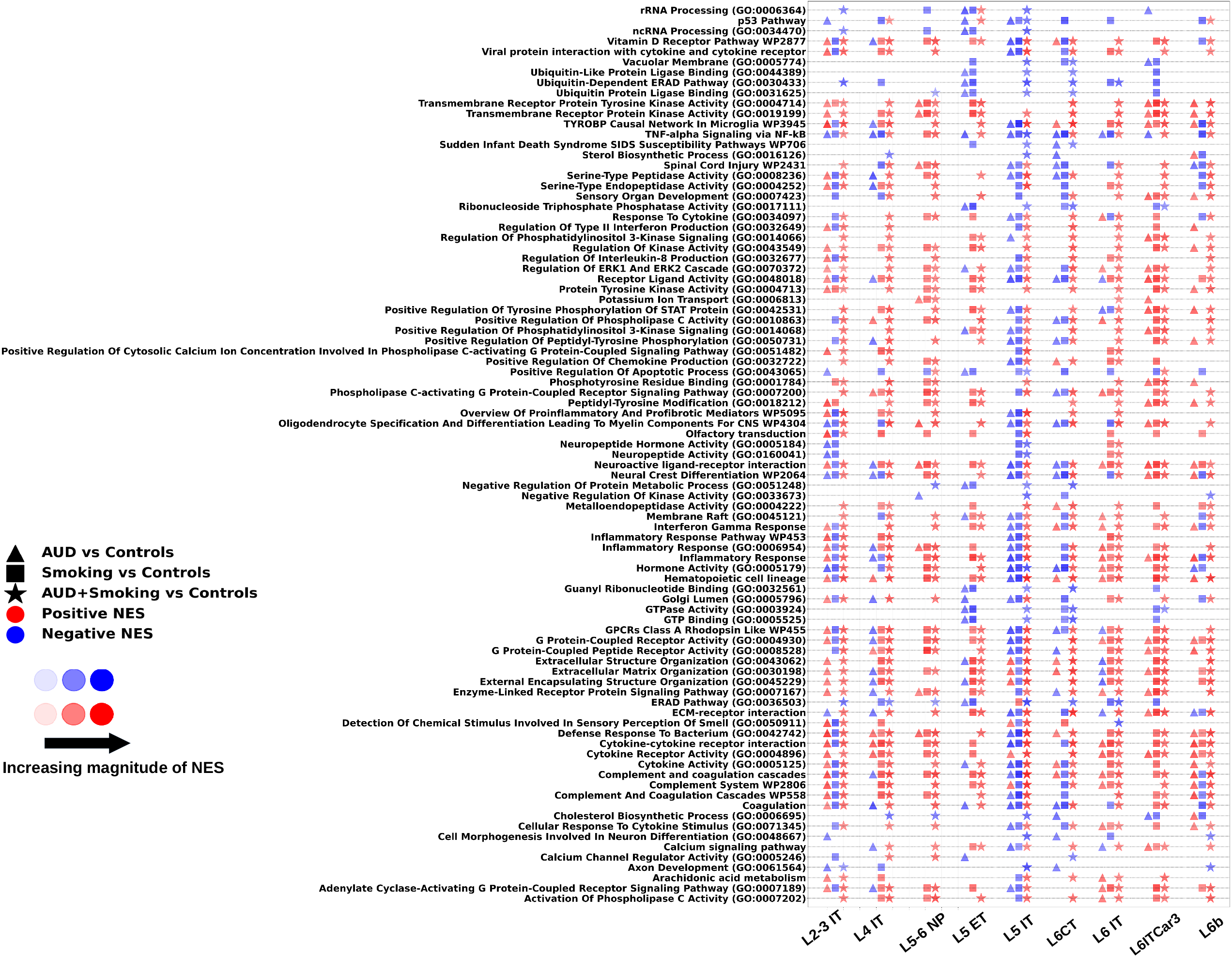
Excitatory neurons’ pathways that are differentially regulated in more cell types in AUD+Smoking vs Controls than AUD vs Controls or Smoking vs Controls. Triangle = AUD vs Controls; Square = Smoking vs Controls; Star = AUD+Smoking vs Controls. Red = positive NES; Blue = negative NES. Symbol size reflects the magnitude of NES.

Complement and coagulation pathways exhibited one of the clearest expansions. Complement system, complement and coagulation cascades, and coagulation gene sets were positively enriched across nearly all excitatory subtypes in AUD+Smoking, whereas they were generally limited to one or two subtypes in AUD and sparse or absent in Smoking alone. The combined condition therefore recruited complement signaling across cortical layers rather than confining it to a small subset of excitatory neurons.

Ubiquitin-proteasome and protein-quality-control pathways followed a similar pattern. Ubiquitin-like protein ligase binding, ubiquitin-protein ligase binding, ubiquitin-dependent ERAD, and related ER-associated degradation programs expanded from limited subtype involvement in the single conditions to L4 IT, L5-6 NP, L5 IT, L6 CT, L6 IT, L6 IT Car3, L6b, and other populations. Thus, proteostatic pathway activation became a broad excitatory-neuron feature in the combined condition.

GPCR signaling showed some of the broadest cell-type expansion. Class A rhodopsin-like GPCRs, G protein-coupled receptor activity, peptide-receptor activity, phospholipase C-coupled signaling, and adenylate cyclase-coupled signaling were positively enriched across most excitatory subtypes in AUD+Smoking. In AUD alone these pathways involved a smaller set of subtypes, and in Smoking alone they were sparse or occasionally negative. Both major GPCR second-messenger arms were therefore broadly engaged when AUD and smoking co-occurred, Figure-2.

Kinase cascades also broadened substantially, including ERK1/2 regulation, general kinase activity, receptor and non-receptor tyrosine-kinase activity, PI3K signaling, and STAT-related phosphorylation. Most were restricted to L2-3 IT or a few additional subtypes in AUD and largely absent in Smoking, but expanded to a majority of excitatory subtypes in AUD+Smoking. Extracellular-matrix organization, extracellular-structure organization, ECM-receptor interaction, and receptor-ligand pathways similarly expanded across most excitatory populations. The pan-excitatory perturbation of ECM pathways could reflect remodeling of the perineuronal and perisynaptic microenvironment, potentially destabilizing synaptic connections across cortical layers (Figure-2).

The expansion was not limited to pathway presence; directionality also became more coordinated. For GPCR and kinase programs, the AUD+Smoking comparison was predominantly positive across subtypes, whereas Smoking alone occasionally produced negative enrichment and AUD alone was more restricted. Calcium-channel regulation, potassium transport, cholesterol and sterol biosynthesis, axon development, myelin/oligodendrocyte-associated programs, oligodendrocyte specification/differentiation, proinflammatory/profibrotic mediator gene sets, neuropeptide hormone activity, and neuroactive ligand-receptor signaling similarly broadened from sparse or subtype-restricted enrichment in the single conditions to multiple excitatory populations in AUD+Smoking; neuronal expression of myelin-related genes has been reported during development [58, 59]. p53 and vitamin D receptor signaling also expanded to additional subtypes. Olfactory transduction was an exception, showing broad negative NES in the combined condition.

These observations indicate that the combined condition, AUD+Smoking, expands not only immune programs but also second-messenger, ion-homeostatic, lipid, developmental, and extracellular-matrix responses across excitatory-neuron classes. Across the neuronal analyses, the combined comparison repeatedly converted sparse single-condition enrichment into multi-subtype engagement, supporting cell-type expansion as the organizing feature of the transcriptional response across the human PFC.

### Inhibitory Neurons’ Responses in AUD with Smoking

Inhibitory neurons showed a parallel but distinct expansion of pathway engagement in AUD+Smoking (Figure-3). A defining feature was recruitment of translational and post-transcriptional regulation. tRNA modification and aminoacylation, miRNA-related programs, mRNA processing, and RNA-splicing pathways were sparse in the single-condition comparisons but expanded across Pax6, Pvalb, Sst, Sst Chodl, Vip, Lamp5, Lamp5 Lhx6, Sncg, and Chandelier populations in the combined condition. Spliceosome and transesterification-related splicing gene sets therefore became broadly engaged across interneuron classes, a pattern less prominent in excitatory neurons.

**Figure-3:**
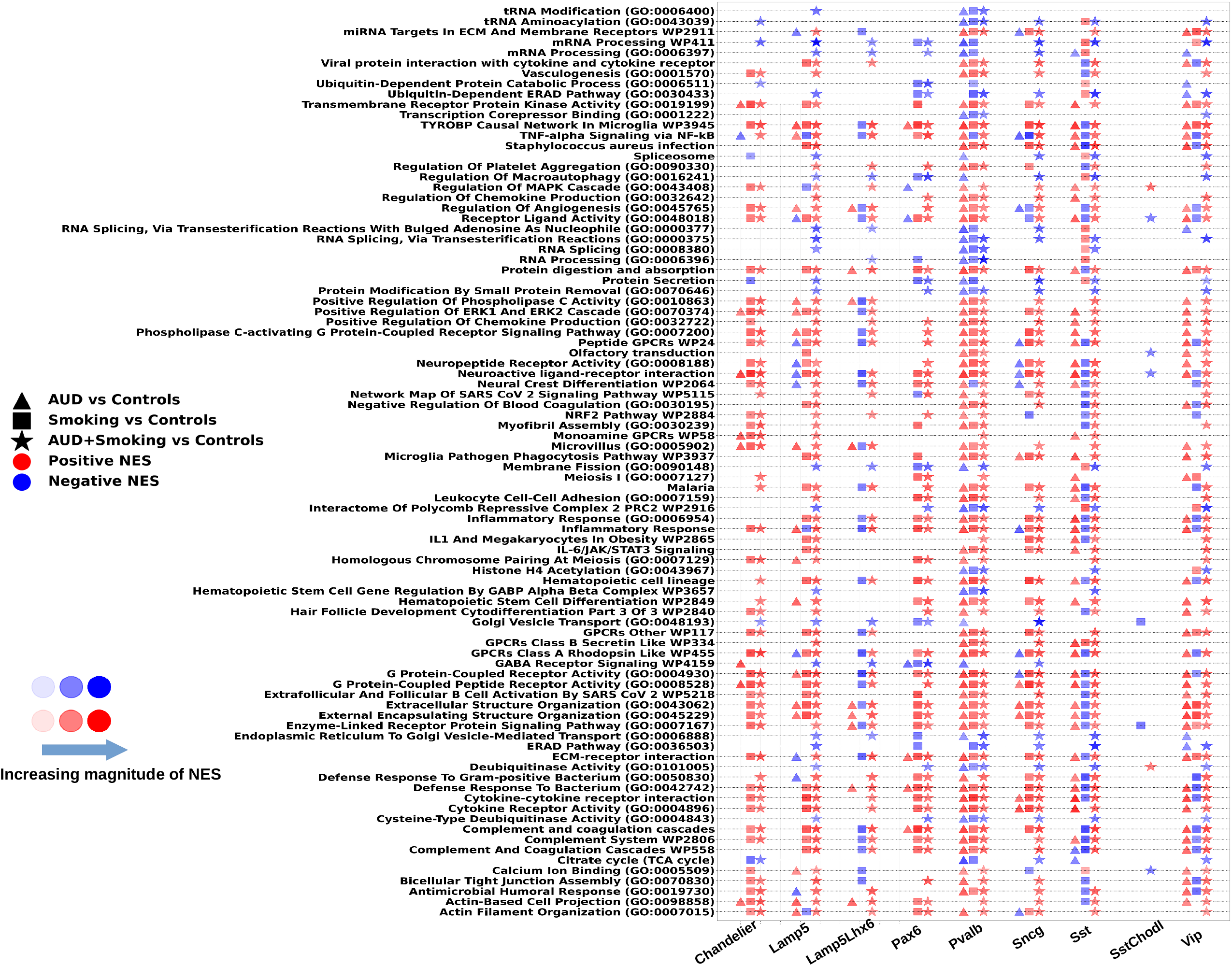
Inhibitory neurons’ pathways that are differentially regulated in more cell types in AUD+Smoking vs Controls than AUD vs Controls or Smoking vs Controls. Triangle = AUD vs Controls; Square = Smoking vs Controls; Star = AUD+Smoking vs Controls. Red = positive NES; Blue = negative NES. Symbol size reflects the magnitude of NES.

Inflammatory pathways also expanded. TNF-alpha/NF-kB and TYROBP signaling, general inflammatory response, viral protein-cytokine interaction, cytokine-receptor pathways, and defense-response programs involved more inhibitory subtypes in AUD+Smoking than in either condition alone. The combined condition additionally recruited IL-6/JAK/STAT3 signaling and related inflammatory gene sets across several subtypes. MAPK regulation and chemokine-production pathways broadened in parallel, linking intracellular kinase cascades with immune-mediator production.

Proteostatic pathways showed a distinctive bidirectional response. Ubiquitin-dependent protein catabolism, ubiquitin-dependent ERAD, and related degradation programs expanded across inhibitory subtypes, but deubiquitinase and cysteine-type deubiquitinase activities also became broadly enriched. This simultaneous recruitment of ubiquitination and deubiquitination distinguishes the inhibitory response from the more unidirectional proteostatic activation seen in excitatory neurons and indicates increased regulation of ubiquitin turnover.

GPCR signaling involved a wider receptor repertoire than in excitatory neurons. In addition to Class A rhodopsin-like and peptide GPCR pathways, Class B secretin-like receptors, monoamine GPCRs, other GPCR classes, and GABA-receptor signaling expanded in the combined condition. Monoamine GPCR enrichment, which was limited to fewer subtypes with Smoking alone, extended across Pvalb, Sncg, Sst, Vip, and other inhibitory populations, indicating broad perturbation of neuromodulatory signaling.

Complement and coagulation, antimicrobial humoral response, cytokine activity, cytokine-receptor interaction, and defense-response pathways were positively enriched across more inhibitory populations. TCA/citrate-cycle pathways, calcium binding, regulation of angiogenesis, bicellular tight-junction assembly, actin organization, membrane fission, microvillus programs, transcription-corepressor binding, macroautophagy, and platelet-aggregation regulation also broadened. Hematopoietic cell-lineage and stem-cell differentiation gene sets, although not indicating literal hematopoietic conversion, reflected shared immune/developmental transcriptional modules and were largely absent from the single-condition comparisons. Importantly, the inhibitory response differed from the excitatory response in both content and subtype distribution: RNA-splicing and translational programs were more prominent, while monoamine, GABA, secretin-like, and peptide GPCR families produced a wider receptor-level signature, together with bidirectional ubiquitin-control signatures.

### Unique Neuronal Effects of Smoking

To delineate smoking-specific effects, we identified pathways significant in Smoking versus controls but not in AUD versus controls, Figure-4 (a), (b). The smoking-specific excitatory-neuron landscape was large and overwhelmingly repressive. Negative NES extended across mitochondrial bioenergetics, protein quality control, RNA processing, cell-survival, and synaptic pathways. Mitochondrial Complex I/OXPHOS, mitochondrial intermembrane-space, TCA-cycle, mitophagy, V-type ATPase, and proton-transport pathways were repressed across several excitatory subtypes, indicating broad impairment of oxidative phosphorylation and proton-gradient maintenance.

**Figure-4:**
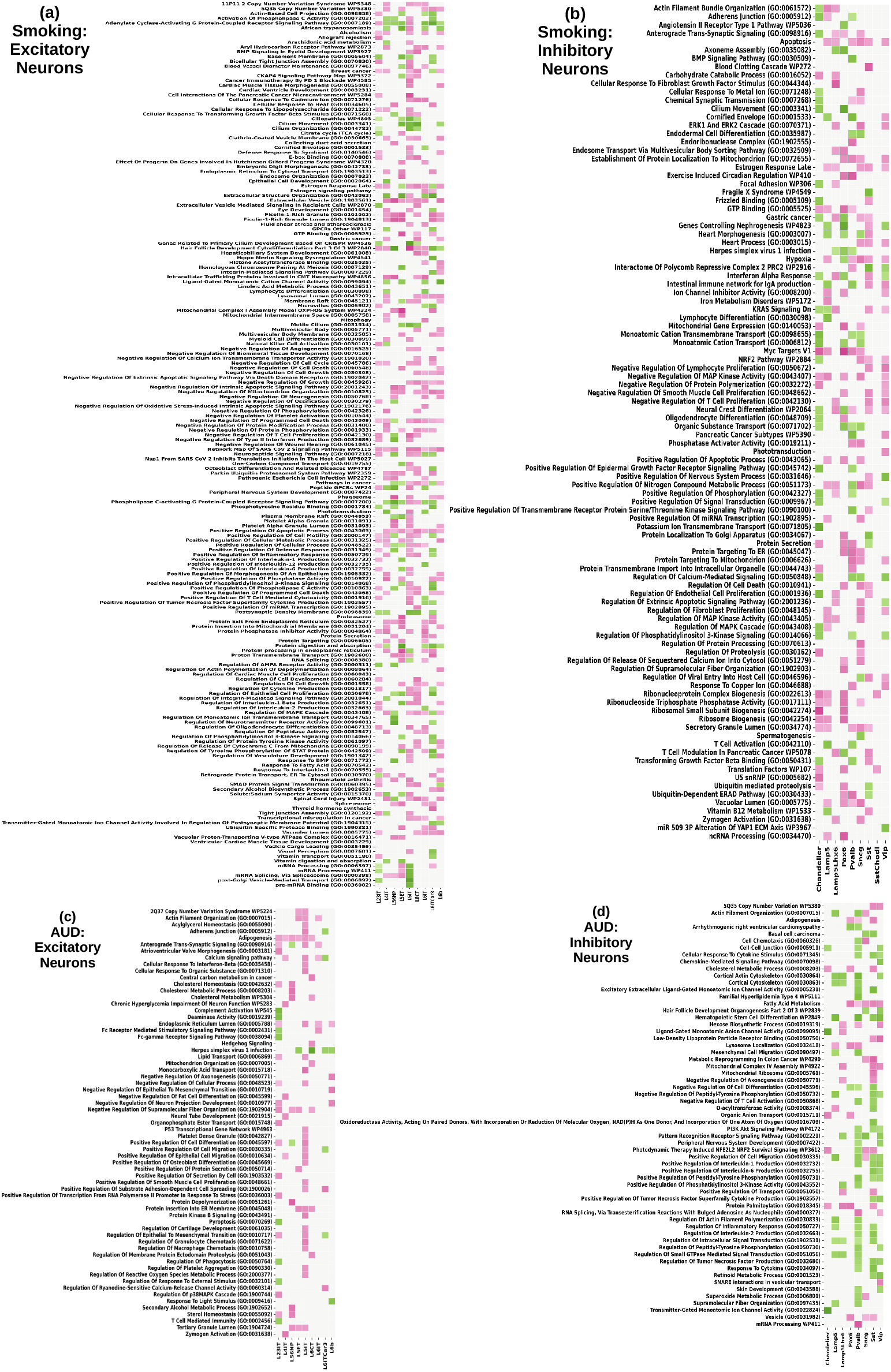
Unique pathways enriched in Smoking vs Controls (a, b in excitatory and inhibitory neurons respectively) and AUD vs Controls (c, d in excitatory and inhibitory neurons respectively).

Proteasome, ER protein processing, ubiquitin-dependent ERAD, mitochondrial-membrane protein insertion, retrograde ER-to-cytosol transport, and protein exit from the ER were repressed. Spliceosome, mRNA processing, mRNA splicing, and pre-mRNA-binding programs also showed negative NES, particularly in deep-layer populations. Multiple anti-apoptotic and cell-survival pathways were suppressed, together with negative regulators of programmed cell death and oxidative-stress-related apoptotic regulation. Synaptic repression included neuropeptide signaling, postsynaptic-density components, AMPA- and broader neurotransmitter-receptor regulation, and transmitter-gated ion-channel activity. Ubiquitin-specific protease binding and peripheral nervous system development were also repressed. These convergent negative signatures indicate that Smoking preferentially suppresses bioenergetic, proteostatic, post-transcriptional, survival, and synaptic functions in excitatory neurons.

Despite this dominant repression, selected deep-layer excitatory populations, particularly L5 ET and L6 IT Car3, showed positive enrichment of phospholipase C- and adenylate cyclase-coupled GPCR signaling, peptide GPCRs, extracellular-structure organization, plasma-membrane raft and integrin-mediated signaling, and inflammatory programs involving IL-1, IL-6, IL-12, and TNF-related cytokine production.

Smoking-specific inhibitory-neuron effects were more heterogeneous. Synaptic transmission and GTP-binding pathways showed mixed positive and negative enrichment across Chandelier, Lamp5, Pax6, Pvalb, Sncg, Sst, and Vip populations. Developmental and structural programs, including BMP signaling, axoneme/cilium pathways, focal and adherens junctions, actin organization, actin-bundle organization, and heart- and nephrogenesis-labeled gene sets, were subtype-specific. Interferon, hypoxia, PRC2, NRF2, ERK, circadian, and oligodendrocyte-differentiation pathways were also selectively enriched. Thus, Smoking showed a much more consistent negative NES profile in excitatory neurons while producing mixed remodeling in inhibitory neurons.

### Unique Neuronal Effects of AUD

AUD-specific pathways were defined as those significant in AUD versus controls but not in Smoking versus controls, Figure-4 (c), (d). Compared with Smoking, the AUD-specific excitatory landscape contained fewer pathways and showed a more bidirectional pattern. Positive NES was concentrated in L5 ET, L6 IT Car3, and selected superficial populations, whereas negative NES occurred across L5-6 NP, L5 IT, L6 CT, and other subtypes. Lipid and sterol metabolism were prominent negative signatures, including cholesterol metabolism/homeostasis, sterol homeostasis, and secondary-alcohol metabolic programs.

In contrast, calcium signaling, Hedgehog signaling, p53 networks, complement activation, Fc-receptor signaling, protein-kinase B signaling, adipogenesis, acylglycerol homeostasis, and other metabolic/developmental programs were enriched in selected excitatory subtypes. AUD-specific excitatory-neuron effects also included Fc-gamma receptor signaling, adipogenic and acylglycerol-homeostasis programs, and multiple pathways regulating cell projection, adhesion, and differentiation. AUD also engaged cell-survival, cell-death, and immune-recruitment pathways. Pyroptosis was enriched together with programs regulating neuron-projection development, fiber organization, cell differentiation, granulocyte and macrophage chemotaxis, platelet aggregation, phagocytosis, T-cell-mediated immunity, and membrane-protein ectodomain proteolysis. This yielded a mixed metabolic-neuroimmune excitatory signature rather than the predominantly repressive pattern seen with Smoking.

The AUD-specific inhibitory-neuron response was dominated by positive NES across multiple subtypes. IL-1, IL-6, IL-2, TNF, inflammatory-response, pattern-recognition receptor, PI3K-Akt, and tyrosine-phosphorylation pathways were broadly engaged, particularly in Pvalb, Sncg, Sst, and Vip neurons. Additional enrichment involved hematopoietic/developmental programs, peripheral nervous system development, NFE2L2/NRF2 survival signaling, cortical cytoskeleton and actin polymerization, SNARE-mediated vesicular transport, ligand-gated ion-channel signaling, fatty-acid and cholesterol metabolism, retinoid metabolism, hexose biosynthesis, protein palmitoylation, mitochondrial Complex IV assembly and mitochondrial-ribosome pathways, and RNA splicing and processing. Selected negative NES involved regulation of axonogenesis, cell differentiation, peptidyl-tyrosine phosphorylation, and T-cell activation. Together, AUD produced broad inflammatory, metabolic, structural, mitochondrial, and post-transcriptional activation in inhibitory neurons and a more heterogeneous excitatory response than Smoking.

### Vascular Perturbations of AUD and Smoking

Vascular cells showed a strong endothelial-VLMC dichotomy that was most evident in AUD+Smoking (Figure-5). Endothelial cells displayed predominantly negative NES, whereas VLMCs showed broad positive enrichment across many pathway classes. In endothelial cells, repression centered on mRNA processing and splicing, mRNA catabolism, translational initiation, transcriptional elongation, rRNA binding, and post-Golgi transport. These RNA-processing deficits were strongest in the combined condition, with some also present in smoking alone.

**Figure-5:**
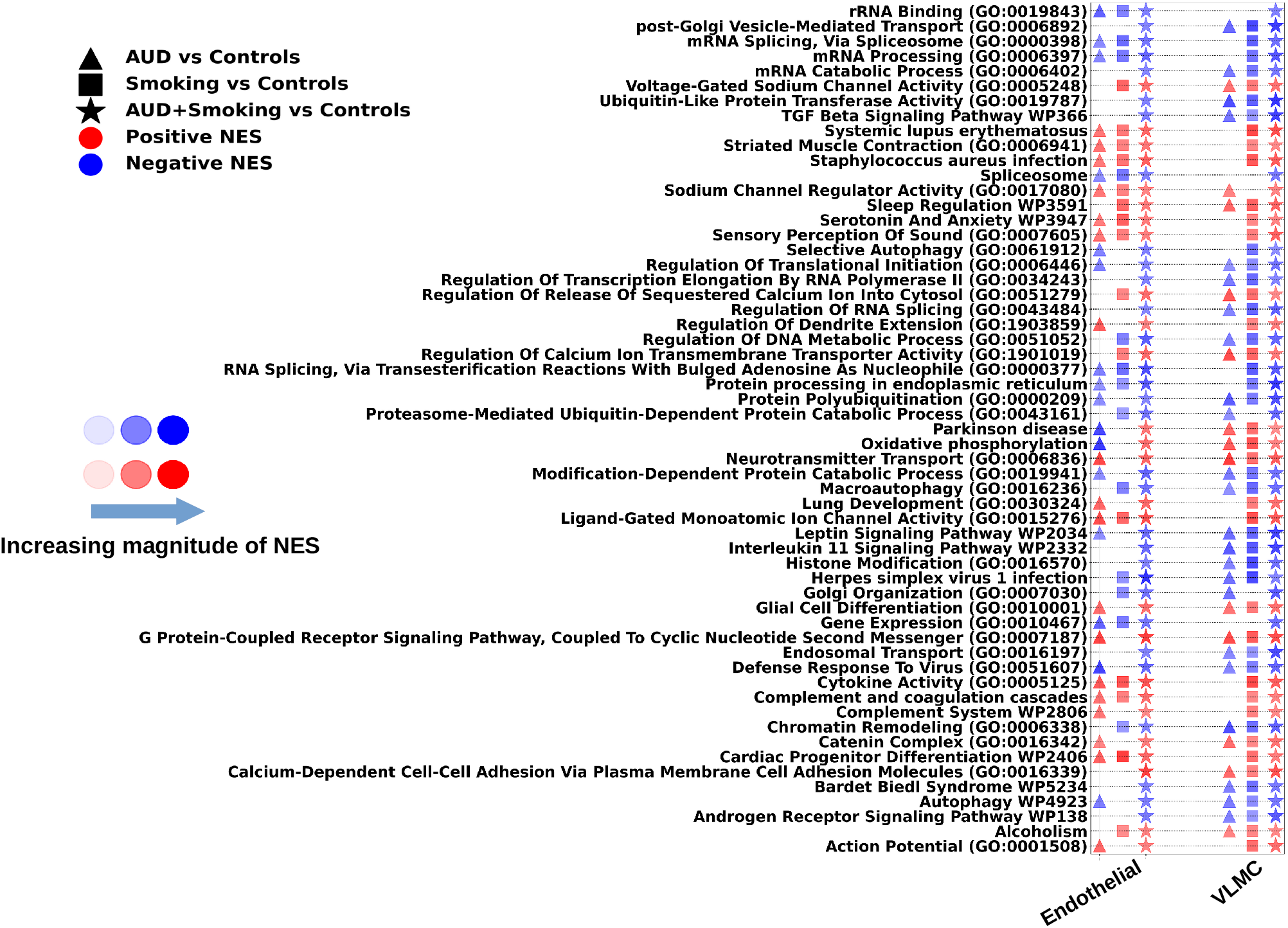
Vascular cells’ pathways that are differentially regulated in more cell types in AUD+Smoking vs Controls than AUD vs Controls or Smoking vs Controls. Triangle = AUD vs Controls; Square = Smoking vs Controls; Star = AUD+Smoking vs Controls. Red = positive NES; Blue = negative NES. Symbol size reflects the magnitude of NES.

Proteostatic and bioenergetic pathways showed the same endothelial direction. Proteasome-mediated and ubiquitin-dependent protein catabolism, protein polyubiquitination, ER protein processing, and ubiquitin-transferase activity were negatively enriched, together with oxidative phosphorylation, macroautophagy, selective autophagy, and related mitochondrial/autophagy programs. Thus, AUD+Smoking concurrently repressed RNA handling, protein turnover, and energy-related functions in endothelial cells.

VLMCs showed the opposite pattern, with positive enrichment of signaling, adhesion, autophagy, calcium-dependent cell-cell adhesion, and metabolic programs. Despite this divergence, complement and coagulation, complement system, cytokine activity, and antiviral/defense pathways were activated in both endothelial cells and VLMCs in AUD+Smoking. This pan-vascular complement engagement paralleled the broad complement expansion observed in excitatory and inhibitory neurons.

Additional vascular perturbations included cyclic-nucleotide GPCR signaling, endosomal and Golgi transport, chromatin remodeling, leptin and IL-11 signaling, TGF-beta signaling, histone modification, DNA metabolic regulation, neurotransmitter transport, and glial-differentiation programs, with several of these pathways particularly evident in VLMCs. Ion-channel and electrophysiological changes included voltage-gated and other sodium-channel pathways, ligand-gated ion channels, and calcium release from intracellular stores. Sleep regulation, serotonin/anxiety-related signaling, and metabolic-inflammatory pathways were also enriched. Although some pathway labels are not vascular-specific, they reflect shared signaling modules within endothelial cells and VLMCs. Overall, the combined condition showed the broadest pathway engagement, reinforcing the endothelial–VLMC divergence across RNA processing, proteostasis, signaling, and inflammatory functions.

### Glial Perturbations of AUD and Smoking

Across astrocytes, microglia, OPCs, and oligodendrocytes, AUD+Smoking engaged pathways in more glial populations than either condition alone (Figure-6). The dominant feature was a marked divergence between microglia and the other three glial cell types. Astrocytes, OPCs, and oligodendrocytes showed predominantly positive NES, whereas microglia showed selective repression of core functional programs. TYROBP causal-network and microglial-phagocytosis pathways were negatively enriched in microglia while showing positive or less-repressed responses in surrounding glia. This pattern paralleled the DEG analysis, in which microglia accounted for 58.1% of repressed DEGs in the combined condition.

**Figure-6:**
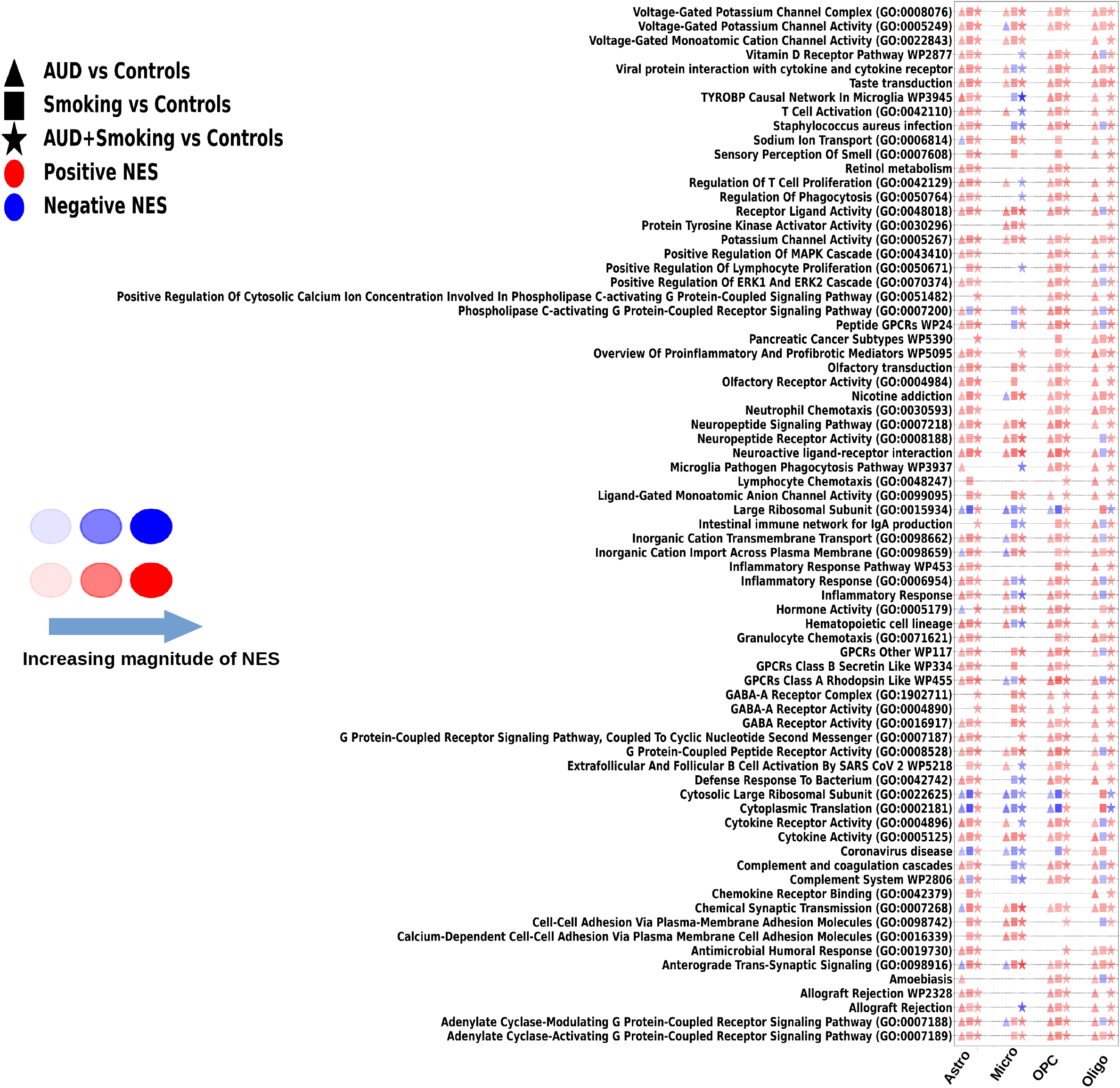
Glial cell-type pathways differentially regulated in more cell types in AUD+Smoking vs Controls than AUD vs Controls or Smoking vs Controls. Triangle = AUD vs Controls; Square = Smoking vs Controls; Star = AUD+Smoking vs Controls. Red = positive NES; Blue = negative NES. Symbol size reflects the magnitude of NES.

Translational machinery reinforced this asymmetry. Large ribosomal-subunit, cytosolic ribosomal-subunit, and cytoplasmic-translation pathways were repressed in microglia but positively enriched in astrocytes, OPCs, and oligodendrocytes. The selective translational repression in microglia occurred together with reduced phagocytic/TYROBP programs, suggesting that the microglial response differed fundamentally from the broad activation observed in other glial cells.

Neuroinflammatory and complement pathways showed some of the widest glial expansion. Inflammatory response, complement and coagulation, complement system, cytokine activity/receptor signaling, and defense-response pathways were positively enriched across astrocytes, OPCs, and oligodendrocytes in AUD+Smoking, with several also activated by AUD alone in astrocytes. Microglia, however, showed mixed or negative NES for several of the same programs. Chemotaxis and immune-cell recruitment pathways, including neutrophil, lymphocyte, and granulocyte chemotaxis, were also positively enriched across non-microglial glia. Additional adaptive-immune-associated gene sets were activated in astrocytes and OPCs while showing weaker or opposite responses in microglia.

GPCR signaling showed extensive glial expansion. Class A and Class B GPCRs, peptide GPCRs, cyclic-nucleotide signaling, phospholipase C-coupled signaling, and adenylate cyclase-coupled pathways were positively enriched across astrocytes, OPCs, and oligodendrocytes. MAPK/ERK and kinase cascades also broadened. GABA-A receptor activity and GABA-receptor-complex pathways were particularly prominent in astrocytes and OPCs, while potassium-, sodium-, voltage-gated cation-, and other ion-transport pathways showed broad positive NES outside microglia, consistent with altered extracellular ion regulation and neuronal excitability. T-cell and lymphocyte proliferation/activation were enriched in astrocytes and OPCs but weaker or negative in microglia.

Neuropeptide and neuroactive ligand-receptor signaling, chemical synaptic transmission and sensory-signaling gene sets, hormone activity, retinol metabolism, vitamin D receptor signaling, profibrotic mediator pathways, extracellular-matrix and calcium-dependent adhesion, and chemokine-receptor binding were also broadly engaged. Some infection- and cancer-labeled pathways were driven by shared immune or ECM genes rather than evidence of those diseases.

Overall, the combined condition produced coordinated inflammatory, GPCR, kinase, ion-channel, and adhesion activation in astrocytes, OPCs, and oligodendrocytes while selectively suppressing microglial phagocytic, TYROBP, and translational programs. This dissociation creates a pronounced asymmetry in the cortical neuroimmune response and distinguishes the combined state from either AUD or Smoking alone.

## Discussion

The present findings show that co-occurring AUD and smoking amplify pathway dysregulation in the human PFC by expanding affected programs across neuronal subtypes. This widespread disruption directly provides a potential molecular basis for evidence that their combined effects on cognition exceed either exposure alone [21, 60, 61]. Neuroimaging studies similarly associate concurrent smoking with greater cortical metabolic abnormalities, gray-matter loss, and impaired recovery of cortical perfusion in AUD [30–32]. Smoking is also associated with poorer neurocognitive recovery during sustained alcohol abstinence [62–64].

In excitatory neurons, activation of inflammatory and cytokine, complement, GPCR, kinase, calcium/ion-channel, proteostatic, and extracellular-matrix pathways expanded from limited subtype involvement in either condition alone to nearly all excitatory populations in combined AUD and smoking. This pattern is consistent with experimental evidence that combined alcohol and tobacco-smoke exposure increases frontal cortical IL-1β and TNF-α beyond either exposure alone [65]. Complement pathways contribute to synaptic pruning and neurodegeneration [66], while the extracellular matrix regulates synaptic plasticity and homeostasis [67]. Inhibitory neurons showed a similar expansion of inflammatory, complement, GPCR, MAPK/chemokine, and proteostatic pathways, together with translational regulation, tRNA modification, RNA processing, splicing, and deubiquitination. Because neuronal identity and circuit function depend heavily on alternative splicing and post-transcriptional regulation [68, 69], these alterations have important implications for cortical inhibitory function and excitatory/inhibitory balance and synaptic plasticity across cortical layers.

Smoking alone produced an excitatory-neuron signature dominated by repression of mitochondrial bioenergetics, proteasomal and ER-associated degradation, RNA processing, cell survival, and synaptic pathways. This profile is consistent with evidence that nicotine alters mitochondrial electron transport and oxidative-phosphorylation programs [70] and directly affects brain mitochondrial respiration [71]. AUD, in contrast, produced broader inflammatory and metabolic activation. The combined condition therefore integrates distinct molecular burdens from each exposure while extending their effects across substantially more cortical neuronal populations.

The vascular findings extend evidence that chronic alcohol exposure and smoking disrupt cerebrovascular function. Alcohol compromises blood-brain barrier integrity through endothelial oxidative stress, tight-junction degradation, and inflammatory signaling involving TLR4, ERK1/2, and NF-κB [72–74], whereas smoking induces endothelial dysfunction through reactive oxygen species, inflammatory cytokines, and matrix metalloproteinases [75–77]. At single-cell resolution, we identify an endothelial–VLMC dichotomy, with repression of RNA processing, proteostatic, and bioenergetic pathways in endothelial cells contrasting with broad VLMC activation. Because VLMCs provide structural support, produce extracellular-matrix components, and contribute to perivascular clearance [78, 79], this opposing response suggests disrupted neurovascular homeostasis within the neurovascular unit. Similar endothelial repression and VLMC activation have been described in Alzheimer’s disease [80, 81].

Despite this divergence, endothelial cells and VLMCs both showed activation of complement and coagulation pathways in combined AUD and smoking. Smoking alone activated complement signaling in VLMCs, whereas the combined condition extended this response across both vascular compartments. This shared complement engagement is relevant because complement-mediated mechanisms contribute to synapse elimination [66] and blood-brain barrier disruption [82], linking the neuronal and vascular abnormalities observed here.

A prior analysis of this cohort, limited to glial cells and AUD, identified inflammatory activation and loss of homeostatic programs in microglia, together with inflammatory astrocyte states [34]. Our analysis extends these findings to smoking and co-occurring AUD and smoking across all PFC cell types. In the combined condition, inflammatory activation expanded across astrocytes, OPCs, and oligodendrocytes, whereas microglia showed repression of TYROBP signaling, phagocytosis, and translation, consistent with an exhaustion-like state. This pattern is reminiscent of microglial dystrophy or exhaustion-like states associated with astrocytic inflammatory activation in aging and Alzheimer’s disease [83–85]. Binge-like alcohol exposure induces microglial dystrophy in rats [86], and alcohol-induced neurodegeneration can combine microglial activation with impaired phagocytic function [87]. Together, these observations support that microglial dysfunction is associated with increased inflammatory activation of surrounding glia and reactive gliosis in combined AUD and smoking.

The contrasting responses of microglia and astrocytes are particularly notable. Activated microglia can induce neurotoxic reactive astrocytes through IL-1α, TNF, and C1q signaling, increasing complement C3 and reducing astrocytic support of synapse formation [88]. Here, astrocytes show activation of complement, cytokine, and inflammatory pathways while these programs are repressed in microglia, suggesting redistribution of inflammatory signaling across glia. Concurrent perturbation of potassium-channel pathways in astrocytes, OPCs, and oligodendrocytes [89, 90] and GABA-receptor pathways in astrocytes and OPCs [91] further implicates glial regulation of extracellular ion homeostasis, gliotransmission, and cortical excitability.

Overall, co-occurring AUD and smoking produce a cortical molecular state broader and qualitatively different from either condition alone across multiple compartments. Across neuronal, glial, and vascular compartments, the most consistent feature is expansion of molecular dysfunction across cell types, with complement signaling emerging as a prominent point of convergence. The microglial exhaustion-like signature, together with inflammatory responses in other glial populations, is consistent with a maladaptive neuroimmune environment associated with prolonged inflammatory stress [86, 88, 92, 93]. These convergent processes provide a plausible molecular substrate for greater neurocognitive impairment in comorbid AUD and smoking [21, 60–63].

These findings provide a rationale for studying therapeutic strategies directed at pathways selectively amplified in the combined condition, including inflammatory, GPCR, kinase, and complement signaling. Future epigenomic, spatial-transcriptomic, and proteomic studies should define the regulatory and anatomical organization of these signatures and determine their relationship to disease severity, treatment response, and accelerated cognitive decline in human brain tissue.

## Limitations of the Study

Because this study analyzed postmortem tissue, it cannot distinguish molecular alterations that confer vulnerability to AUD and smoking from those resulting from chronic exposure. Studies of human brain tissue across different stages of disease progression, together with epigenomic and spatial analyses and longitudinal studies of accessible biomarkers, may help distinguish predisposing alterations from the consequences of chronic exposure and define their regulatory and anatomical context.

## Data and Code Availability

Data used in this work is publicly available via GEO (with accession number GSE247416) as a part of the publication [34]. Code used to analyze the data is on the GitHub repository https://github.com/aj95b/AUD_Smoking.

## Acknowledgement

This work was supported by NIH grants AA028982, AA021667, DA056004, DA058399, DA063157. We also thank Anna Warden and R. Dayne Mayfield at the Waggoner Center for Alcohol and Addiction Research for providing details about the dataset and promptly addressing queries.

## Conflict of Interest

Authors declare no conflicts of interest.

## Notes

### Competing Interest Statement

The authors have declared no competing interest.

