## Supplementary Figure-1 for "Alcohol Use Disorder and Smoking-Associated Molecular Alterations in Human Prefrontal Cortex in Single Nucleus (sn) RNA-Seq"

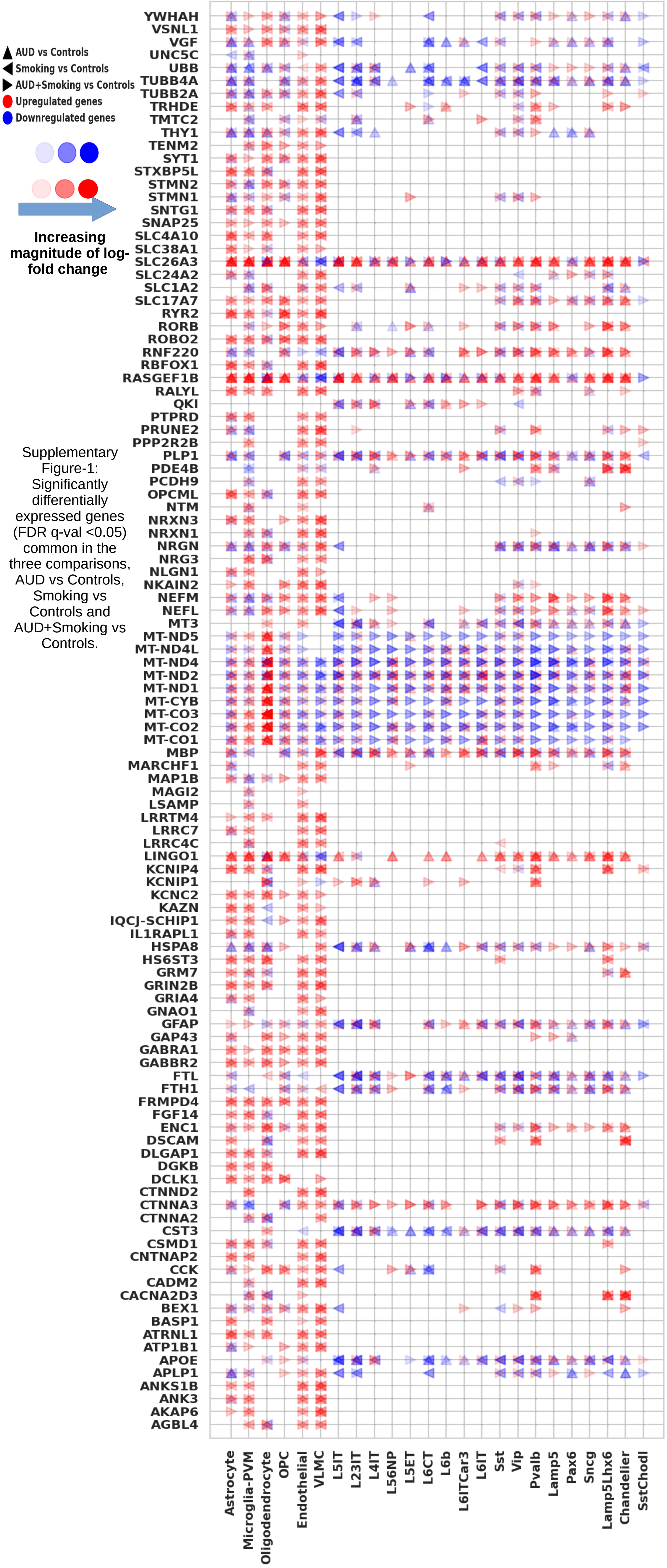


Supplementary Figure-1: Significantly differentially expressed genes (FDR q-val <0.05) common in the three comparisons, AUD vs Controls, Smoking vs Controls and AUD+Smoking vs Controls.
